# Defining Relictual Biodiversity: Conservation Units in Speckled Dace (Cyprinidae: *Rhinichthys osculus*) of the Greater Death Valley Ecosystem

**DOI:** 10.1101/2020.06.09.143339

**Authors:** Steven M. Mussmann, Marlis R. Douglas, David D. Oakey, Michael E. Douglas

## Abstract

The tips in the tree of life serve as foci for conservation and management, yet clear delimitations are masked by inherent variance at the species-population interface. Analyses using thousands of nuclear loci can potentially sort inconsistencies, yet standard categories applied to this parsing are themselves potentially conflicting and/or subjective [e.g., DPS (distinct population segments); DUs (Diagnosable Units-Canada); MUs (management units); SSP (subspecies); Evolutionarily Significant Units (ESUs)]. One potential solution for consistent categorization is to create a comparative framework by accumulating statistical results from independent studies and evaluating congruence among data sets. Our study illustrates this approach in speckled dace (Cyprinidae: *Rhinichthys osculus*) endemic to two basins (Owens and Amargosa) in the Death Valley ecosystem (DVE). These fish persist in the Mojave Desert as isolated Pleistocene-relicts and are of conservation concern, but lack formal taxonomic descriptions/designations. Double-digest RAD (ddRAD) methods identified 14,355 SNP loci across 10 populations (N=140). Species delimitation analyses [multispecies coalescent (MSC) and unsupervised machine learning (UML)] delineated four putative ESUs. *F*_ST_ outlier loci (N=106) were juxtaposed to uncover the potential for localized adaptations. We detected one hybrid population that resulted from upstream reconnection of habitat following contemporary pluvial periods, whereas remaining populations represent relics of ancient tectonism within geographically-isolated springs and groundwater-fed streams. Our study offers three salient conclusions: A blueprint for a multi-faceted delimitation of conservation units; a proposed mechanism by which criteria for intraspecific biodiversity can be potentially standardized; and a strong argument for the proactive management of critically-endangered DVE fishes.

## 1 INTRODUCTION

Species represent the currency of biodiversity, and as such are focal points for conservation and management. Yet, they often lack those distinct demarcations expected of their categorization, and represent instead mere waypoints along evolutionary pathways (Sullivan et al., 2014). Our capacity to discriminate is further confounded by an observed variance in intraspecific diversity that also contributes to the difficulties in unambiguously defining taxonomic units. Furthermore, a multiplicity of species concepts (Zachos, 2018) not only adds to, but also underscores this difficulty. ‘What precisely is a species?’ and ‘How can it be delineated?’ are questions with both philosophical and practical ramifications (de Queiroz, 2007). As a result, the lack of resolution promotes confusion among managers tasked with deciding what should be conserved, and how best to accomplish the task (Douglas, Douglas, Schuett, Porras, & Thomason, 2007; Holycross & Douglas, 2007). It also represents what we now define as the ‘species problem’ (Freudenstein, Broe, Folk, & Sinn, 2017; Garnett & Christidis, 2017).

Taxonomic ranks above the species are commonly agreed upon and represent human constructs arbitrarily delimited by systematists (Coyne & Orr, 2004), with entities shuffled and re-shuffled according to morphological and molecular perspectives (and the limitations thereof). Taxa are assigned to these ranks based upon the knowledge and opinion of practitioners (see for example, Yang et al., 2015). Surprisingly, this approach has also been widely adopted for categorizations below the species rank. This represents an extension of the species-problem, where terminology has not only become confusing but also conflicting. We address these issues below.

### 1.1 A focus on intraspecific categorization

Intraspecific lineages often stem from a single ancestral source, potentially one population or a cluster thereof. Their components are relatively contiguous geographically, albeit with temporal dynamics, such as demographic expansions, contractions, and even potential stasis within refugia as repercussions of climatic fluctuation (Levin, 2019). They have been variously labeled, from more traditional frameworks such as subspecies or ecological races (Braby, Eastwood, & Murray, 2012), through more contemporary concepts including evolutionarily significant units (ESUs) and management units (MUs) (Coates, Byrne, & Moritz, 2018), to those promoted by government regulations, such as distinct population segments (DPS: USFWS & NMFS, 1996) or diagnosable units (DUs: COSEWIC, 2012). The focus for all these categorizations remains unitary: The recognition and conservation of intraspecific diversity.

The emergence of molecular techniques to quantify ‘genetic diversity’ was initially heralded as a potential solution for the demarcation of biodiversity units, but traditional methods provided scant applicability with regard to identifying intraspecific boundaries, and thus served to extend classificatory confusion (Phillimore & Owens, 2006; Zink, 2004). The advent of next-generation sequencing (NGS) techniques has markedly improved our ability to evaluate questions surrounding intraspecific variation, although they too present a double-edged sword: Discrete population-level patterns can indeed be identified, but an over-interpretation of these patterns is a management issue in that populations are often inappropriately categorized. These fears are borne out as well through complex analytical techniques commonly used to delimit species (Campillo, Barley, & Thomson, 2019; Sukumaran & Knowles, 2017). In addition, sophisticated analytical methods are now emerging, such as Machine Learning (ML), that employ pattern recognition as a mechanism to identify biodiversity, and which allow systems to automatically learn and improve from experience without being explicitly so programmed.

We outline in this manuscript a multifaceted approach for the delimitation of species and intraspecific diversities. We first: (1) define the questions to be addressed, (2) detail different analytical approaches that quantify inter- and intra-specific diversity using NGS data, (3) place detected patterns within a spatial and temporal framework so as to understand underlying evolutionary and ecological drivers, and (4) combine our data with those available from prior publications and grey literature reports to evaluate diversity within the context of adaptive potential and ecological boundaries (Cornetti, Ficetola, Hoban, & Vernesi, 2015; Stanton et al., 2019). We offer this multifaceted approach. (Figure 1) as a potential blueprint from which to delimit biologically distinct entities, and as a mechanism to classify intraspecific biodiversity via standardized criteria for recognition as conservation units.

**Figure 1:**
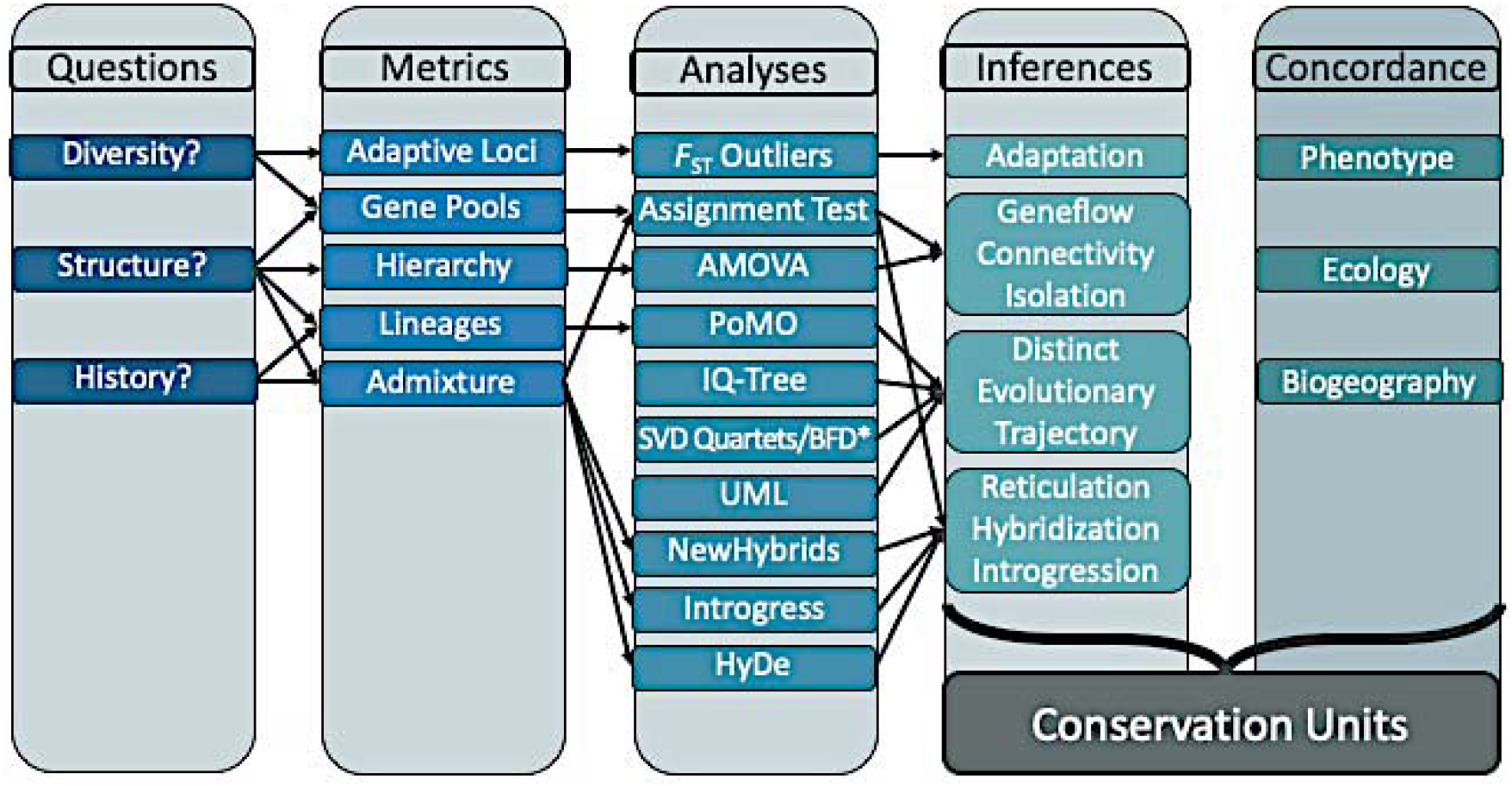
Conceptual workflow to delineate conservation units at tips in the Tree of Life. A standardized mechanism that applies molecular data to categorize inter- and intra-specific diversity should address three **Questions** about patterns in genetic data that must be integrated with other available data (**Concordance**). **Metrics** depict genetic diversity and are quantified singly or in combination via **Analyses** to derive **Inferences** regarding how ecological and evolutionary processes have shaped these patterns. Genetic results and other public data (e.g., phenotypic, ecological, biogeographical) are combined in a comparative framework so as to depict conservation units.

To illustrate this approach, we offer a case study using an endemic group of desert fishes whose conservation is clearly hampered by taxonomic ambiguity. The advantages of our study are its various levels of complexity, both intra- and interspecific, manifested within a simple spatial design (i.e., populations within isolated habitats), within a region well understood with regards to biogeography and paleohydrology. We use our framework to develop a strong argument for the proactive management of critically endangered fishes in one of the world’s most unique environments, the Death Valley Ecosystem (DVE) of arid Southwestern North America.

### 1.2 Study species and its biogeography

The issues introduced above globally transcend regions, ecosystems and organisms. We focus in this study on a desert ecosystem (Mojave Desert) with aquatic organisms largely restricted to freshwater springs (Craig, Kollaus, Behen, & Bonner, 2016; Devitt, Wright, Cannatella, & Hillis, 2019; Morvan et al., 2013). The latter support a surprisingly high level of species-richness (Jetz, Rahbek, & Colwell, 2004) across springs, as driven by a post-Pleistocene desiccation that effectively eliminated congeners/ competitors (Smith, 1981), thus evoking simplicity within springs. Most are relictual (Grandcolas, Nattier, & Trewick, 2014), with species diversity largely underestimated (Rossini, Fensham, Stewart-Koster, Gotch, & Kennard, 2018), despite the lack of closely related taxa (Minckley, Hendrickson, & Bond, 1986; Smith, Badgley, Eiting, & Larson, 2010).

Our study species is a small cyprinid (speckled dace: *Rhinichthys osculus* = SPD) from the Death Valley ecosystem (DVE) of southwestern Nevada and eastern California (Figure 2). Although broadly distributed through western North America (Furiness, 2012; Oakey, Douglas, & Douglas, 2004; Sada, Britten, & Brussard, 1995), it reaches greatest diversity in the DVE [i.e., five subspecies of ‘special concern’ (Moyle, Quiñones, Katz, & Weaver, 2015), of which only one (*R. o. nevadensis*) is formally described (Deacon & Williams, 1984; Gilbert, 1893; La Rivers, 1962; Williams, Hardy, & Deacon, 1982)]. The unique environmental conditions manifested within the DVE have engendered multiple, allopatric populations distributed across two basins, Owens and Armagosa (Figure 2). These populations lack a formal taxonomic description and this consequently constrains their federal protection to ‘distinct population segments’ (Haig et al., 2006). Given this, we consider these unidentified entities in this study as being OTUs (operational taxonomic units; Sokal & Sneath, 1963).

**Figure 2:**
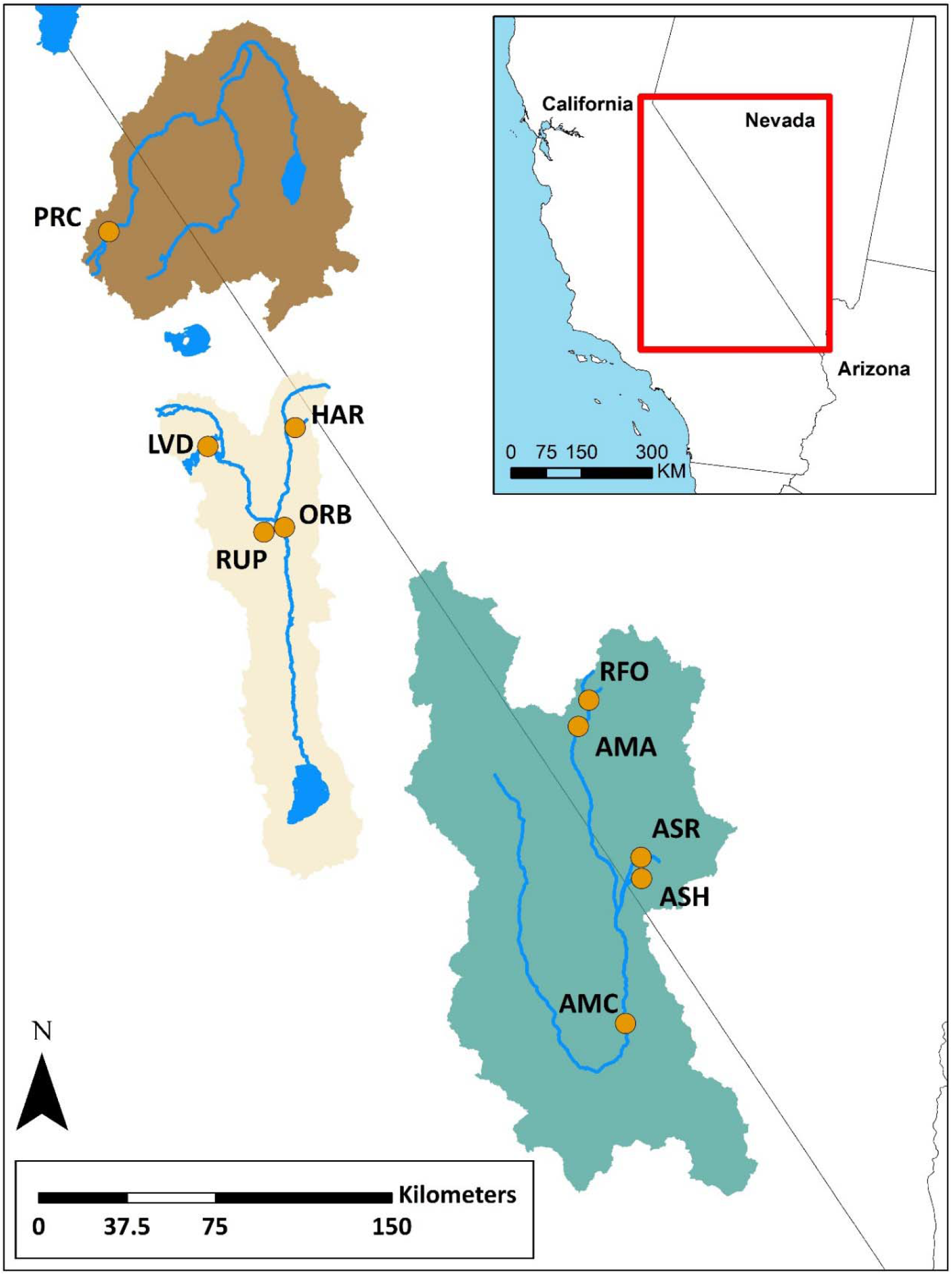
Sampling sites for speckled dace (*Rhinichthys osculus*; SPD) within the Walker (dark brown), Owens (light brown) and Amargosa (green) river basins. SPD localities are: PRC=Walker Sub-basin (*R. o. robustus*); AMA and RFO=Oasis Valley; AMC=Amargosa Canyon; ASH and ASR=Ash Meadows (*R. o. nevadensis*); HAR=Benton Valley; ORB and RUP=Owens Valley; LVD=Long Valley.

While both basins are endorheic (with no connection to other basins), their hydrology differs. The high desert of the Owens Basin exists in the rain shadow of the eastern Sierra Nevada, and accumulates surface water through snowmelt runoff from the surrounding mountains. However, available habitat for SPD has been systematically depleted by aqueducts constructed in 1913 and 1970 to supply the metropolis of Los Angeles. This, in turn, has forced considerable reliance upon local groundwater resources (Danskin, 1998). In contrast, the Amargosa Basin has been historically arid (Belcher, Sweetkind, Hopkins, & Poff, 2019), with intermittent flows in the Amargosa River a residual of rare, large-scale precipitation (Grasso, 1996). Here, SPD persists in ground water seeps (Faunt, Blainey, Hill, & D’Agnese, 2019) that have remained relatively consistent despite overexploitation of local water resources (Robbins, 2017).

This study provides a comprehensive, population-level evaluation of SPD as a case study for conservation unit delineation in a system with confused intraspecific variability. We sequenced thousands of nuclear loci (SNPs) so as to: (1) Gauge gene flow within and among populations; (2) Identify nuclear loci under selection as a proxy for local adaptation; (3) Generate a phylogeny using species tree methods; and (4) Employ MSC and machine learning algorithms to delimit intraspecific diversity. As a means to validate conservation units and underscore management, we then (5) interpreted the genetic data in the context of morphological and ecological descriptions available from published sources. This comparative framework also allowed us to interpret the evolutionary drivers that shaped this unique and endemic biodiversity.

## 2 METHODS

### 2.1 Sampling

The Owens Basin (i.e., Long Valley and Owens River) and the Amargosa Basin (i.e., Ash Meadows and Amargosa River) were our sampling regions. Populations were evaluated from nine locations (Figure 2). These represented five putative OTUs in DVE (Owens: 4 localities, N=50 samples; Amargosa: 5 localities, N=80 samples). Sampling spanned 1989-2017, with Long Valley sampled twice (1989, 2016) and Oasis Valley three times (1993, 2004, 2016). Two closely related taxa served as outgroups: eastern blacknose dace (N=5; *R. atratulus*, Rogue River, MI) and Lahontan speckled dace [N=10; *R. o. robustus*, Poore Creek, Walker sub-basin (PRC, Figure 2)]. Table 1 provides an overview, with additional information on the taxonomic history of these OTUs as detailed in Appendix 1.

**Table 1:**
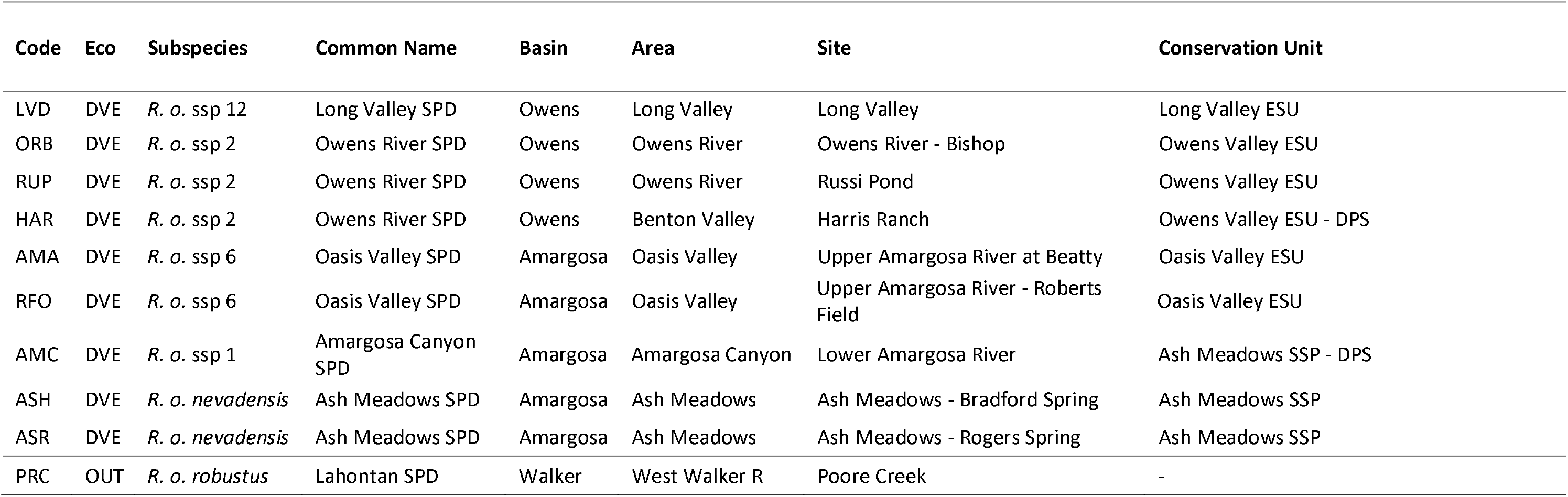
Speckled dace (*Rhinichthys osculus*) sample groups analyzed in this study, including the ingroup from the Death Valley Ecosystem (DVE) and an outgroup from the Lahontan Basin (OUT). Latin trinomials are provided for described taxa. Those lacking formal description are provided subspecies (ssp) numbers reflecting those assigned by fisheries managers for unnamed *R. osculus* taxa in the Great Basin and surrounding areas. Common names, basins, regions within basins (Area) and specific site names (Site) are provided. Conservation units reflect assignments designated by multispecies coalescent and machine learning algorithms (ESU = Evolutionarily Significant Unit; DPS = Distinct Population Segment).

| Code | Eco | Subspecies | Common Name | Basin | Area | Site | Conservation Unit |
| --- | --- | --- | --- | --- | --- | --- | --- |
| LVD | DVE | *R. o. ssp 12* | Long Valley SPD | Owens | Long Valley | Long Valley | Long Valley ESU |
| ORB | DVE | *R. o. ssp 2* | Owens River SPD | Owens | Owens River | Owens River - Bishop | Owens Valley ESU |
| RUP | DVE | *R. o. ssp 2* | Owens River SPD | Owens | Owens River | Russi Pond | Owens Valley ESU |
| HAR | DVE | *R. o. ssp 2* | Owens River SPD | Owens | Benton Valley | Harris Ranch | Owens Valley ESU - DPS |
| AMA | DVE | *R. o. ssp 6* | Oasis Valley SPD | Amargosa | Oasis Valley | Upper Amargosa River at Beatty | Oasis Valley ESU |
| RFO | DVE | *R. o. ssp 6* | Oasis Valley SPD | Amargosa | Oasis Valley | Upper Amargosa River - Roberts Field | Oasis Valley ESU |
| AMC | DVE | *R. o. ssp 1* | Amargosa Canyon SPD | Amargosa | Amargosa Canyon | Lower Amargosa River | Ash Meadows SSP - DPS |
| ASH | DVE | *R. o. nevadensis* | Ash Meadows SPD | Amargosa | Ash Meadows | Ash Meadows - Bradford Spring | Ash Meadows SSP |
| ASR | DVE | *R. o. nevadensis* | Ash Meadows SPD | Amargosa | Ash Meadows | Ash Meadows - Rogers Spring | Ash Meadows SSP |
| PRC | OUT | *R. o. robustus* | Lahontan SPD | Walker | West Walker R | Poore Creek | - |

### 2.2 Data collection and filtering

Whole genomic DNA was extracted using several methods: Gentra Puregene DNA Purification Tissue kit; QIAGEN DNeasy Blood and Tissue Kit; QIAamp Fast DNA Tissue Kit; and CsCl-gradient. Extracted DNA was visualized on 2.0% agarose gels and quantified with a Qubit 2.0 fluorometer (Thermo Fisher Scientific, Inc.). Library preparation followed a double digest Restriction-Site Associated DNA (ddRAD) protocol (Peterson, Weber, Kay, Fisher, & Hoekstra, 2012). Barcoded samples (100 ng DNA each) were pooled in sets of 48 following Illumina adapter ligation, then size-selected at 375-425 bp (Chafin, Martin, Mussmann, Douglas, & Douglas, 2018) using the Pippin Prep System (Sage Science). Size-selected DNA was subjected to 12 cycles of PCR amplification using Phusion high-fidelity DNA polymerase (New England Bioscience) following manufacturer protocols. Subsequent quality checks to confirm successful library amplification were performed via Agilent 2200 TapeStation and qPCR. Final libraries were pooled in sets of three per lane and subjected to 100bp single-end sequencing (Illumina HiSeq 2000, University of Wisconsin Biotechnology Center; and HiSeq 4000, University of Oregon Genomics & Cell Characterization Core Facility).

Libraries were de-multiplexed and filtered for quality using process_radtags (Stacks v1.48; Catchen, Hohenlohe, Bassham, Amores, & Cresko, 2013). All reads with uncalled bases or Phred quality scores < 10 were discarded. Reads with ambiguous barcodes that otherwise passed quality filtering were recovered when possible (= 1 mismatched nucleotide). A clustering threshold of 0.85 (Eaton, 2014) was used for *de novo* assembly of ddRAD loci in PYRAD v3.0.66. Reads with > 4 low quality bases (Phred quality score < 20) were removed. A minimum of 15 reads was required to call a locus for an individual. A filter was applied to remove putative paralogs by discarding loci with heterozygosity > 0.6 and those containing > 10 heterozygous sites. The resulting data were filtered (BCFtools; Li, 2011) as a means of retaining a single biallelic SNP from each locus, as present in at least 33% of individuals (hereafter referred to as ‘SNP-all’). Filtering was designed to minimize potential bias in missing data for ingroup samples, given the unbalanced basin sampling (e.g., Owens N=50 *versus* Amargosa N=80) (Eaton, Spriggs, Park, & Donoghue, 2017; Huang & Knowles, 2016).

### 2.3 Loci under selection

To identify potential for local adaptation, all loci were subjected to *F*_ST_ outlier analysis (i.e., *F*_ST_ outliers indicate loci under selection). BCFtools-filtered SNPs were analyzed in BayeScan v2.1 (Foll & Gaggiotti, 2008), using recommended settings [20 pilot runs (5,000 generations each) followed by 100,000 Markov Chain Monte Carlo (MCMC) generations (including 50,000 burn-in)]. Data were thinned by retaining every 10^th^ sample, equating to 5,000 total MCMC samples. Outlier status was determined by a false discovery rate (FDR) of 0.05.

BayeScan has the lowest Type I and II error rates among comparable software, yet a single outlier-detection method elicits some level of uncertainty (Narum & Hess, 2011). Thus, cross-validation was conducted using the FDIST2 method in Lositan (Antao, Lopes, Lopes, Beja-Pereira, & Luikart, 2008; Beaumont & Nichols, 1996). Conditions included 100,000 total simulations, assuming a ‘neutral’ forced mean *F*_ST_, 95% confidence interval (CI), and FDR of 0.1. Those SNPs deemed to be under positive selection by both BayeScan and Lositan were extracted for downstream analysis (=SNP-select).

### 2.4 Population structure analysis

A Maximum Likelihood (ML) approach (Alexander, Novembre, & Lange, 2009) was utilized to assess population structure in both datasets. A minor allele frequency filter (AdmixPipe v2.0; Mussmann, Douglas, Chafin, & Douglas, 2020) was applied to remove SNPs at a frequency < 0.01. The number of distinct gene pools in the data set was explored in Admixture using clustering (K) values of 1 to 20, each with 20 replicates. Cross-validation (CV) values were calculated following program instructions.

The output was evaluated using a Markov clustering algorithm to identify different modes calculated by Admixture within a single K-value so as to automate the process of summarizing multiple independent Admixture runs (Clumpak; Kopelman, Mayzel, Jakobsson, Rosenberg, & Mayrose, 2015). Major clusters were identified at a similarity threshold of 0.9 and summarized in AdmixPipe. The best interpretation of population structure was the K-value associated with the lowest CV score.

To further assess population structure and distribution of genetic diversity, an analysis of molecular variance (AMOVA; Excoffier & Lischer, 2010) was performed for the full SNP dataset (SNP-all), and again for SNP-select. In both cases, pairwise *F*_ST_ values served to evaluate genetic isolation between basins and among localities within basins. For sites with temporal collections, AMOVA was also used to test for genetic variance among sampling events within localities.

### 2.5 Hybridization

Preliminary evaluations indicated high levels of Admixture at one site, Amargosa Canyon, the most downstream site in the Armagosa River drainage (Figure 2). Three approaches were used to determine whether hybridization occurred, and if so, its relative timing. First, the Bayesian clustering program NewHybrids v1.1 beta 3 (Anderson & Thompson, 2002) was employed to assess Amargosa Canyon samples. SNPs were filtered in GenePopEdit to obtain a set of unlinked loci that maximized *F*_ST_ among populations (Stanley, Jeffery, Wringe, DiBacco, & Bradbury, 2017). The ‘z’ option in NewHybrids was used to assign two ‘pure’ parental gene pools (upstream sites in Armagosa Basin: Oasis Valley and Ash Meadows) to contrast individuals with admixed ancestry (Armagosa Canyon). The probability was then evaluated for classification of samples into each of six categories: Pure Oasis Valley; Pure Ash Meadows; first generation (F1) Oasis Valley by Ash Meadows hybrid; second generation (F2) hybrid; F1 backcross with pure Oasis Valley; or, F1 backcross with pure Ash Meadows. The program was run for 1,000,000 generations of burn-in followed by 3,000,000 generations of data collection.

Our second approach was to calculate a hybrid index (Buerkle, 2005) using the est.h function in the introgress R package (Gompert & Buerkle, 2010). Data were filtered so as to acquire only fixed, unlinked SNPs among parental populations. Interspecific heterozygosity was calculated using the calc.intersp.het function in introgress, and results were visualized using the triangle.plot function.

Finally, HyDe (Blischak, Chifman, Wolfe, & Kubatko, 2018) was used to test whether Amargosa Canyon was of hybrid origin. This method differs from the previous two in that it tests for hybrid lineages by employing phylogenetic invariants that arise under the coalescent model. Bonferroni adjustment (α=0.0045) was used to test for significance, and 500 bootstrap replicates were performed.

### 2.5 Phylogenetic Analyses

To identify distinct evolutionary lineages, species tree methods were used in phylogenetic analyses using all samples (N=135), including the outgroup taxa [*R. atratulus* and *R. o. robustus* (PRC)], but excluding a population from a private pond (RUP) because of its uncertain origin. To account for incomplete lineage sorting, the reversible polymorphism-aware phylogenetic model (PoMo) was applied in IQ-Tree v1.6.9 (Nguyen, Schmidt, von Haeseler, & Minh, 2014; Schrempf, Minh, De Maio, von Haeseler, & Kosiol, 2016). This method allows polymorphic states to occur within populations, rather than following the traditional assumption in DNA substitution models that taxa are fixed for a specific nucleotide at a given locus. To model genetic drift, a virtual population size of 19 was assumed. Mutations followed a GTR substitution model, and rate heterogeneity was modeled using four categories. An ultrafast bootstrap algorithm (Hoang, Chernomor, von Haeseler, Minh, & Le, 2018) performed 1,000 bootstrap replicates.

SVDQuartets (Chifman & Kubatko, 2014; Chifman & Kubatko, 2015), as implemented in PAUP* (Swofford, 2003), was used to construct a phylogeny using a multispecies coalescent approach. This method evaluates all combinations of four populations (quartets) at each locus, then calculates a single value decomposition (SVD) score (Golub & Van Loan, 1996) for each possible quartet tree. The topology with the lowest SVD score is selected as the true quartet topology. The full species tree is then assembled using a quartet assembly algorithm (Reaz, Bayzid, & Rahman, 2014). Exhaustive quartet sampling was conducted, and significance was assessed using 100 bootstrap replicates.

### 2.6 Conservation unit delimitation

Two approaches were employed to determine the number of discrete conservation units in the Death Valley region: Multispecies coalescent (MSC) methods and unsupervised machine learning (UML) algorithms. Machine learning algorithms have been proposed as alternatives to MSC methods, which seemingly over-split taxa under certain conditions (Barley, Brown, & Thomson, 2018; Leaché, Zhu, Rannala, & Yang, 2018), and parse populations rather than speciation events (Sukumaran & Knowles, 2017). This stems from the many assumptions implicit to MSC, such as random mating, neutral markers, a lack of post-speciation gene flow, and no within-locus recombination or linkage disequilibrium (Degnan & Rosenberg, 2009). Furthermore, UML methods reduce subjectivity in that they do not rely upon user-defined models (Derkarabetian, Castillo, Koo, Ovchinnikov, & Hedin, 2019).

A MSC-based method (BFD*: Leaché et al. 2014) was first employed to test if observed genetic diversity could be divided into discrete, well-supported units (i.e., subspecies, populations, or geographic subdivisions). Data were filtered in the PHRYNOMICS R package (Leaché, Banbury, Felsenstein, de Oca, & Stamatakis, 2015) to remove invariant sites, non-binary SNPs, and loci appearing in >95% of individuals. However, these conditions demanded impractical compute times for each model, so we randomly subsampled the filtered data to retain 200 SNPs and five samples from each of eight sample sites (N=40). These sites represented the most recent temporal collections of genetic clusters identified by Admixture.

We estimated the prior value for the population mutation rate (Θ) in BFD* using the mean pairwise sequence divergence (7.99 × 10^−3^) between *R. osculus* and its sister taxon *R. cataractae* (Longnose Dace). This value was set as the mean of a gamma-distributed prior. The lineage birth rate (λ) of the Yule model was fixed using PYULE (https://github.com/joaks1/pyule). A λ-value of 181.49 assumed tree height as one-half of the maximum observed pairwise sequence divergence. Path sampling was set to 48 steps of 500,000 MCMC generations, with 100,000 discarded as burn-in. Bayes factors (BF) were calculated from normalized marginal likelihoods (Leaché et al., 2014) using the current taxonomy (i.e., five subspecies) as a reference point. Models were evaluated for statistical significance via BF (Kass & Raftery, 1995).

Five UML algorithms were also applied (Derkarabetian et al., 2019), to include: Discriminant analysis of principal components (DAPC: Jombart & Ahmed, 2011); Random Forest (RF) methods (Liaw & Wiener, 2002) with classical (RF cMDS) and isotonic (RF isoMDS) multidimensional scaling; t-distributed stochastic neighbor embedding (t-SNE: van der Maaten & Hinton, 2008); and variational autoencoder (VAE: Kingma & Welling, 2013). For each RF and t-SNE method, the optimal number of clusters (K) was determined by partitioning around medoids (PAM) with gap statistic using k-means clustering (Kassambara & Mundt, 2019) and via hierarchical clustering (Scrucca, Fop, Murphy, & Raftery, 2016). These algorithms were applied to all naturally occurring SPD populations (N=130), using the 200 SNPs previously sampled for BFD* (i.e., SPD from Russi Pond (RUP) and *R. atratulus* samples were excluded).

## 3 Results

### 3.1 Alignment, filtering, and loci under selection

PYRAD recovered 15,020 loci (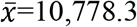; =2,288.3) present in at least 33% of the ingroup samples. Mean sequencing depth per locus was 62.08x (*σ*=21.17x). After filtering by BCFtools, 14,355 loci were retained. Missing data (28.12% total) ranged from 8.42% to 70.96% (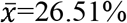; *σ*=13.57%) within individual samples.

BayeScan recovered 210 loci under positive selection, whereas Lositan found 632. We cross-referenced them to acquire a consensus of 106 loci (the “SNP-select” dataset) representing a subsample of the 14,355 locus “SNP-all” data.

### 3.2 Population structure

Genetic diversity within the Owens and Amargosa basins was best represented by seven genetic clusters representing SNP-all (Figure 3A) and SNP-select data (Figure 3B), with Lahontan SPD (PRC) forming an additional 8^th^ group. In (A), all proposed subspecies (Table 1) were recovered as unique populations, with two subspecies being further subdivided: (1) Owens River was separated into an upstream cluster in Benton Valley (HAR) and a downstream cluster with locations near Bishop, CA (RUP/ ORB); and (2) Oasis Valley was also parsed into upstream (RFO) and downstream (AMA) groups. The SNP-select results (Figure 3B) showed a similar trend but with slightly greater admixture, mostly among Amargosa Basin populations. Unique genetic signatures corresponded to the same eight clusters recovered from the SNP-all dataset (Figure 3A).

**Figure 3:**
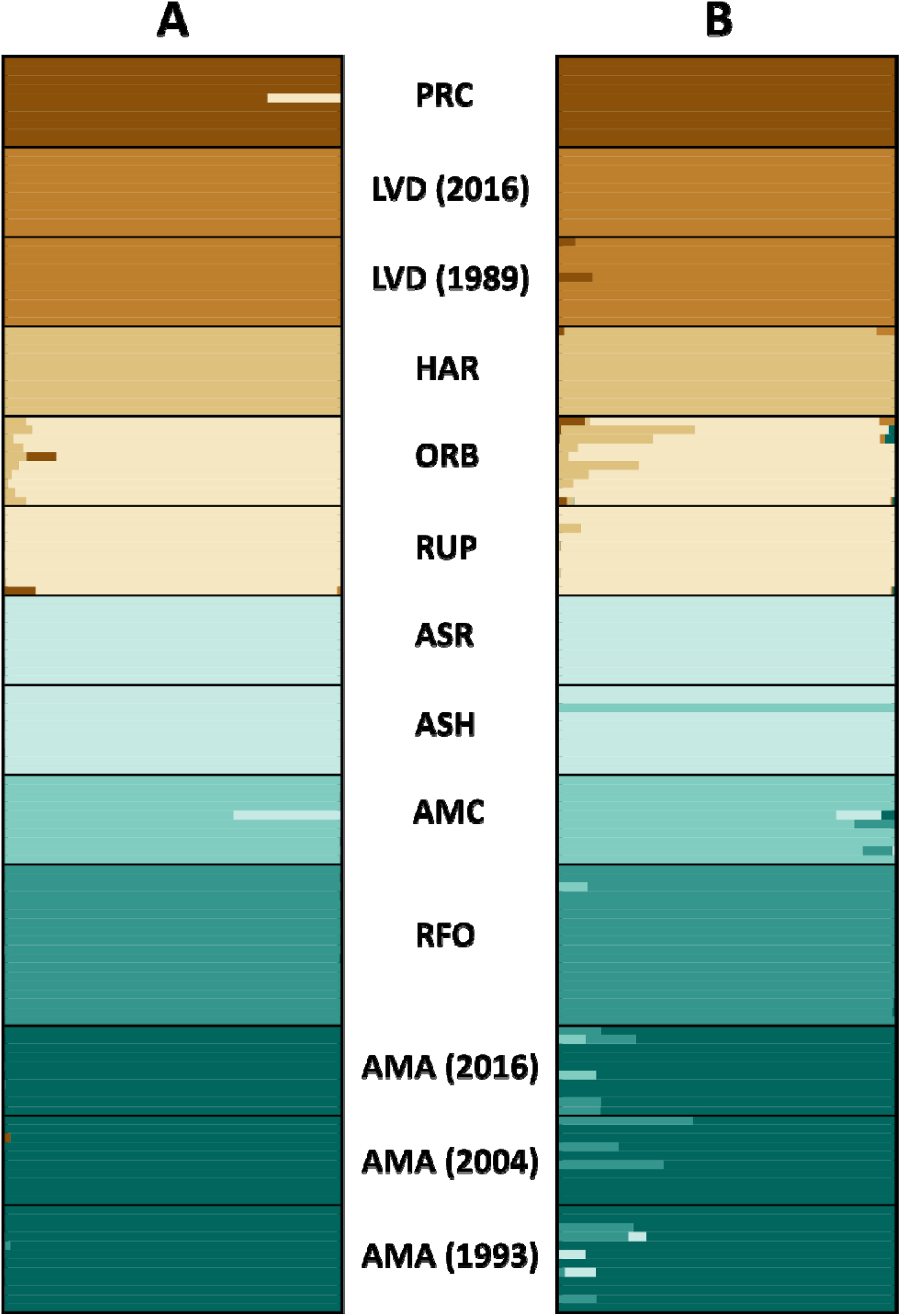
Results of Admixture analyses showing among speckled dace (*Rhinichthys osculus*; SPD) population structure. (A) Analysis of the full SNP dataset (SNP-all; 14,355 loci). (B) Analysis of SNPs determined to be under selection by *F*_ST_ outlier analyses (SNP-select; 106 loci). Localities are: PRC=Walker Sub-basin (*R. o. robustus*); AMA and RFO=Oasis Valley; AMC=Amargosa Canyon; ASH and ASR=Ash Meadows (*R. o. osculus nevadensis*); HAR=Benton Valley; ORB and RUP=Owens Valley; LVD=Long Valley. Numbers next to locality names represent year of collection.

AMOVA results for both datasets revealed high genetic divergence among localities (SNP-all *F*_ST_=0.50; SNP-select *F*_ST_=0.96; p<0.001). Genetic divergence among Admixture-defined clusters was also high (SNP-all *F*_CT_=0.48; SNP-select *F*_CT_=0.94; p<0.001), but with variability among localities reduced within clusters (SNP-all *F*_SC_=0.04; SNP-select *F*_SC_=0.28). The proportion of genetic variance distributed among Admixture-identified clusters was greatest for SNP-select (SNP-all =48.02%; SNP-select =93.82%) whereas for SNP-all it was within sampling localities (SNP-all =49.93%; SNP-select =4.43%). The proportion of variance distributed among localities within clusters was very low for both (SNP-all =2.05%; SNP-select =1.75%) indicating that stochastic temporal sampling variance is unlikely the cause of the observed strong genetic differences among localities.

Pairwise *F*_ST_ values revealed significant isolation among sampling localities for both datasets (Table 2). All pairwise comparisons for SNP-all were significant, save a comparison of two Owens Valley populations (ORB/ RUP), and two temporal comparisons within Oasis Valley (AMA-1993 *versus* 2004 and 2016). The greatest pairwise values were between Long Valley (LVD) *versus* other localities (Mean *F*_ST_=0.706; range=0.623-0.784). The second greatest comparison (Mean *F*_ST_=0.540; range=0.318-0.783) also involved an Owens Basin population, the upstream location in Benton Valley (HAR). Similar trends were observed for SNP-select, with the most notable exception being a lack of significant pairwise *F*_ST_ values when comparing temporal sampling events within-populations, which strongly suggests that stochastic temporal sampling is not driving the differences among sites. Pairwise *F*_ST_ values were greater overall, with two Owens Basin locations again reflecting elevated divergence [Long Valley (LVD): Mean *F*_ST_=0.950; range=0.849-0.992) and Benton Valley (HAR): Mean *F*_ST_=0.925; range=0.685-0.994, respectively].

**Table 2:** Pairwise *F*_ST_ values calculated via AMOVA in Arlequin for Amargosa and Owens River basin speckled dace (*Rhinichthys osculus*). Cells shaded in green represent comparisons among Amargosa Basin localities, whereas those in brown compare localities within the Owens River Basin. Values below diagonal were calculated for the full dataset of 14,355 loci (=SNP-all). Values above the diagonal were calculated from 106 loci determined to be under selection (=SNP-select). All *F*_ST_ values are significant at Bonferroni-adjusted p<0.0038, save those in bold. Sites for which there were multiple sampling events (AMA, LVD) also reflect the year of collection. SPD locations are: PRC=Walker Subbasin (*R. o. robustus*); AMA and RFO=Oasis Valley; AMC=Amargosa Canyon; ASH and ASR=Ash Meadows (*R. o. nevadensis*); HAR=Benton Valley; ORB and RUP=Owens Valley; LVD=Long Valley. Numbers next to sampling locality names represent years during which repeated collections occurred.

|  | PRC | LVD (2016) | LVD (1989) | ORB | RUP | HAR | AMA (1993) | AMA (2004) | AMA (2016) | RFO | ASH | ASR | AMC |
| --- | --- | --- | --- | --- | --- | --- | --- | --- | --- | --- | --- | --- | --- |
| PRC | * | 0.851 | 0.926 | 0.887 | 0.913 | 0.947 | 0.965 | 0.968 | 0.976 | 0.973 | 0.999 | 0.998 | 0.992 |
| LVD (2016) | 0.669 | * | **0.101** | 0.849 | 0.880 | 0.888 | 0.958 | 0.960 | 0.965 | 0.968 | 0.985 | 0.985 | 0.980 |
| LVD (1989) | 0.646 | 0.031 | * | 0.887 | 0.905 | 0.929 | 0.964 | 0.968 | 0.973 | 0.973 | 0.992 | 0.992 | 0.988 |
| ORB | 0.334 | 0.655 | 0.623 | * | 0.343 | 0.685 | 0.942 | 0.941 | 0.948 | 0.957 | 0.970 | 0.970 | 0.968 |
| RUP | 0.392 | 0.650 | 0.627 | **0.000** | * | 0.801 | 0.945 | 0.942 | 0.949 | 0.960 | 0.973 | 0.972 | 0.968 |
| HAR | 0.551 | 0.783 | 0.764 | 0.333 | 0.318 | * | 0.969 | 0.972 | 0.977 | 0.977 | 0.994 | 0.993 | 0.989 |
| AMA (1993) | 0.485 | 0.699 | 0.672 | 0.332 | 0.343 | 0.502 | * | **-0.004** | **-0.022** | 0.608 | 0.817 | 0.792 | 0.773 |
| AMA (2004) | 0.500 | 0.748 | 0.721 | 0.365 | 0.335 | 0.556 | **0.002** | * | **0.019** | 0.612 | 0.854 | 0.845 | 0.796 |
| AMA (2016) | 0.464 | 0.717 | 0.682 | 0.331 | 0.301 | 0.510 | **-0.032** | 0.015 | * | 0.647 | 0.869 | 0.860 | 0.817 |
| RFO | 0.478 | 0.684 | 0.658 | 0.330 | 0.350 | 0.492 | 0.197 | 0.189 | 0.162 | * | 0.866 | 0.850 | 0.786 |
| ASH | 0.579 | 0.784 | 0.763 | 0.418 | 0.407 | 0.614 | 0.396 | 0.440 | 0.395 | 0.392 | * | 0.946 | 0.938 |
| ASR | 0.552 | 0.776 | 0.750 | 0.408 | 0.397 | 0.594 | 0.398 | 0.441 | 0.406 | 0.391 | 0.199 | * | 0.915 |
| AMC | 0.432 | 0.701 | 0.665 | 0.286 | 0.267 | 0.479 | 0.237 | 0.262 | 0.243 | 0.240 | 0.303 | 0.296 | * |

### 3.3 Hybridization

A hybrid index was calculated for crosses between Ash Meadows (ASH) and the two Oasis Valley populations (AMA and RFO). Data represented fixed differences between the two [i.e., 61 SNPs (AMA x ASH: Figure 4A) and 83 SNPs (RFO x ASH: Figure 4B)]. The genomic composition of each Amargosa Canyon sample is an approximate 50/50 representation of each parent, with hybridization ranging from recent (F2) to historic based upon observed interspecific heterozygosity (i.e., interspecific heterozygosity < 0.5). NewHybrids demonstrated a similar trend using the 390-locus dataset filtered in GenePopEdit (Figure 4C). All Amargosa Canyon samples were classified with high probability as F2 hybrids (P>0.97).

**Figure 4:**
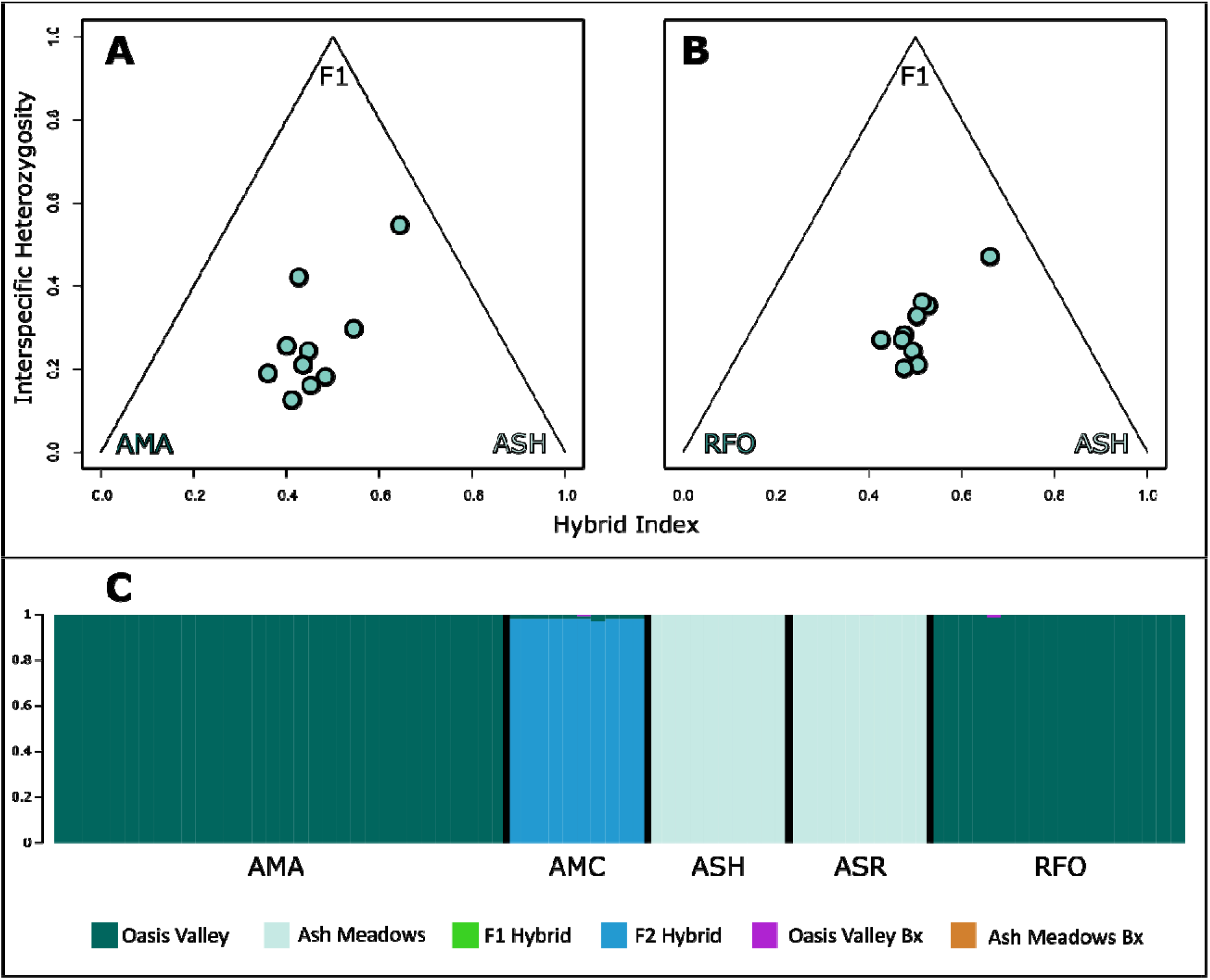
Results of hybrid tests for Amargosa Canyon (AMC) speckled dace (SPD; *Rhinichthys osculus*). (A) and (B) represent hybrid indices plotted against interspecific heterozygosity for two Oasis Valley populations, AMA and RFO. Ash Meadows (ASH and ASR) was used as the second parental population. SNPs were filtered to find fixed differences among populations (AMA x ASH= 61 SNPs; RFO x ASH = 83 SNPs; = SNP-select). (C) The same individuals were classified in each of six categories using a 390-locus dataset. These categories included pure Oasis Valley SPD; pure Ash Meadows SPD; F1 hybrids; F2 hybrids; Oasis Valley backcross (Bx); and Ash Meadows backcross (Bx). All AMC SPD were classified as F2 hybrids with high probability (P>0.97).

Results of HyDe (Table 3) were congruent with the above, again indicating a hybrid origin for Amargosa Canyon. Both Ash Meadows localities (ASH and ASR) were significant [P<0.0045; with high bootstrap support (=97.8-100)] when compared with Oasis Valley SPD at Beatty (AMA). Only Ash Meadows samples from Bradford Spring (ASH) yielded a significant result with moderate bootstrap support when compared against Oasis Valley samples from Roberts Field (RFO).

**Table 3:**
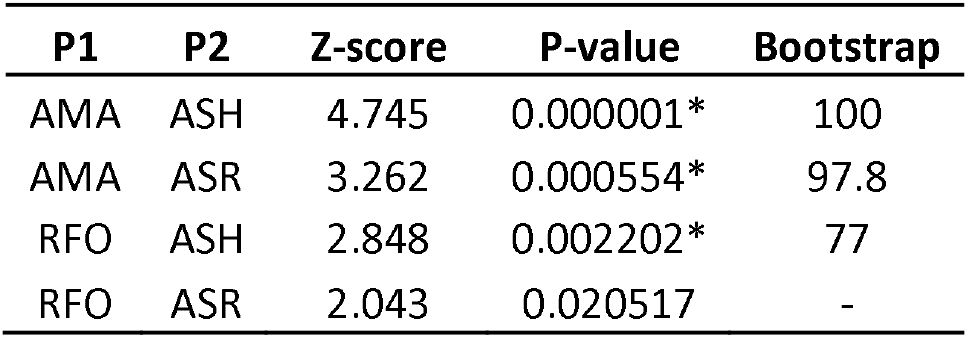
Test for a hybrid origin of Amargosa Canyon speckled dace using HyDe and 14,355 loci (=SNP-all). Comparisons were conducted for both Ash Meadows localities (ASH=Bradford Spring; ASR=Rogers Spring) with Oasis Valley samples (AMA=Amargosa River near Beatty, NV; RFO = Roberts Field). Significant Bonferroni-adjusted p-values (P<0.0045) indicated by asterisks (*). Five hundred bootstrap replicates were performed for each test.

### 3.4 Phylogenetic analyses

PoMo and SVDQuartets results converged on the same tree topology. While the five Amargosa Basin sites formed a clade, Long Valley SPD (LVD) forced a paraphyletic Owens Basin group (Figure 5). Most nodes were well supported, with the greatest uncertainty surrounding placement of the hybrid Amargosa Canyon (AMC) lineage as sister to the Ash Meadows clade (ASH and ASR; SVDQuartets bootstrap = 62). The other uncertainty concerned the relationship between the downstream Owens River (ORB) and the upstream Benton Valley (HAR) populations, which together formed a moderately supported clade in both methods (SVDQuartets bootstrap=87; PoMo bootstrap=74).

**Figure 5:**
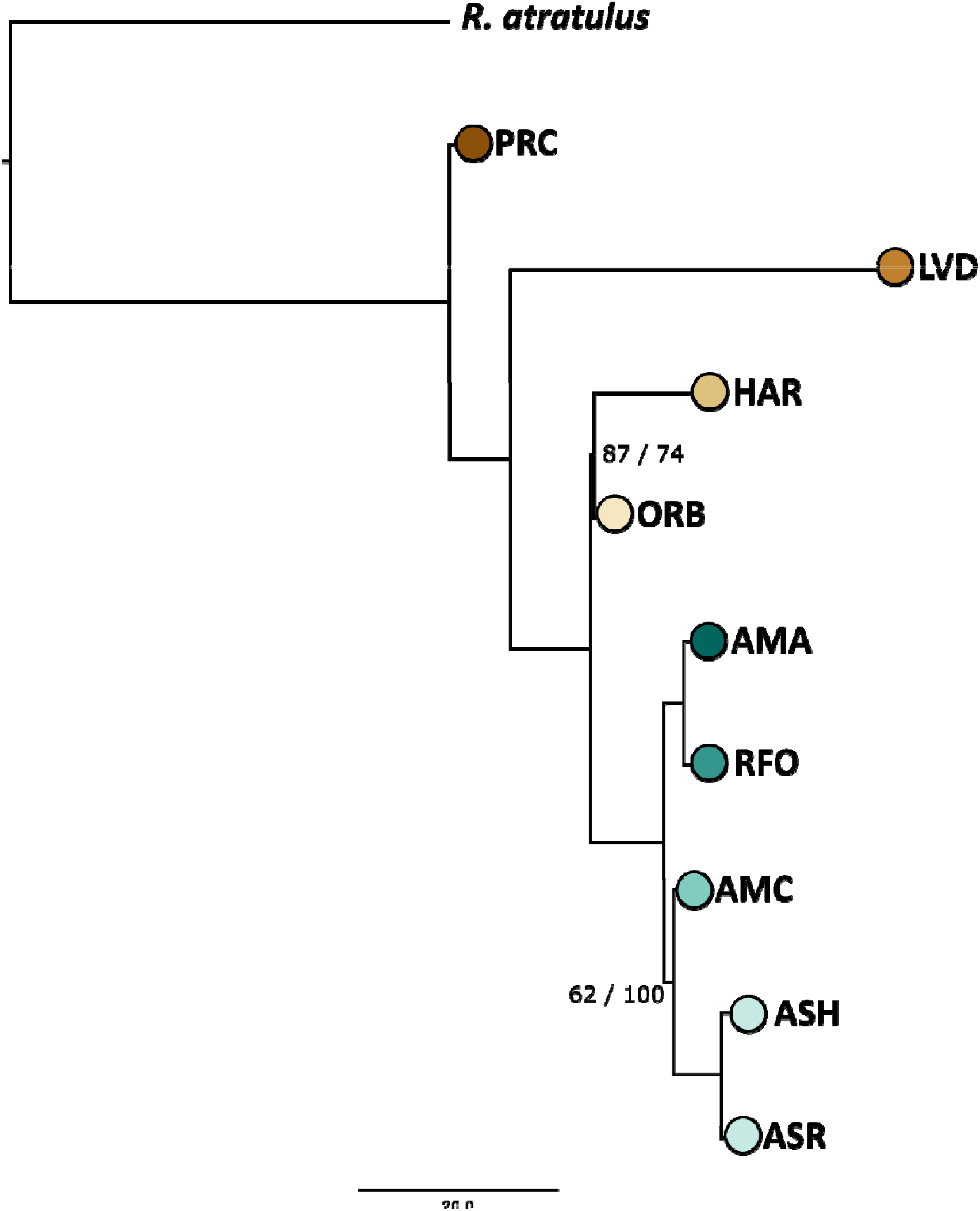
Phylogenetic tree of Owens and Amargosa basin speckled dace (SPD: *Rhinichthys osculus*) sampling localities. Analysis employed the SVDQuartets multispecies coalescent approach (MSC) in PAUP*, and polymorphism-aware models (PoMo) in IQ-TREE. Branch lengths from IQ-TREE are displayed. Bootstrap support values from SVDQuartets (left) and PoMo (right) are displayed only for nodes with support <100. Eastern blacknose dace (*R. atratulus*) was used as outgroup. SPD localities are: PRC=Walker Sub-basin (*R. o. robustus*); AMA and RFO=Oasis Valley; AMC=Amargosa Canyon; ASH and ASR=Ash Meadows (*R. o. nevadensis*); HAR=Benton Valley; ORB and RUP=Owens Valley; LVD=Long Valley.

### 3.5 Bayes Factor Delimitation

The BFD* analysis (Figure 6) decisively split the dataset into eight unique lineages, with seven distributed between the Owens and Amargosa basins (BF=501.85). In this regard, BF≥10 is considered very strong support (Kass & Raftery, 1995). The seven groups correspond to those populations previously identified by Admixture [i.e., five subspecies, with the two Owens River (HAR and ORB) and the two Oasis Valley (AMA and RFO) populations each being segregated]. The next two most highly ranked models collapsed subspecies within the Amargosa Basin: The first collapsed Oasis Valley populations into a single entity (BF=397.58), while the second grouped Ash Meadows with Amargosa Canyon (BF=228.43).

**Figure 6:**
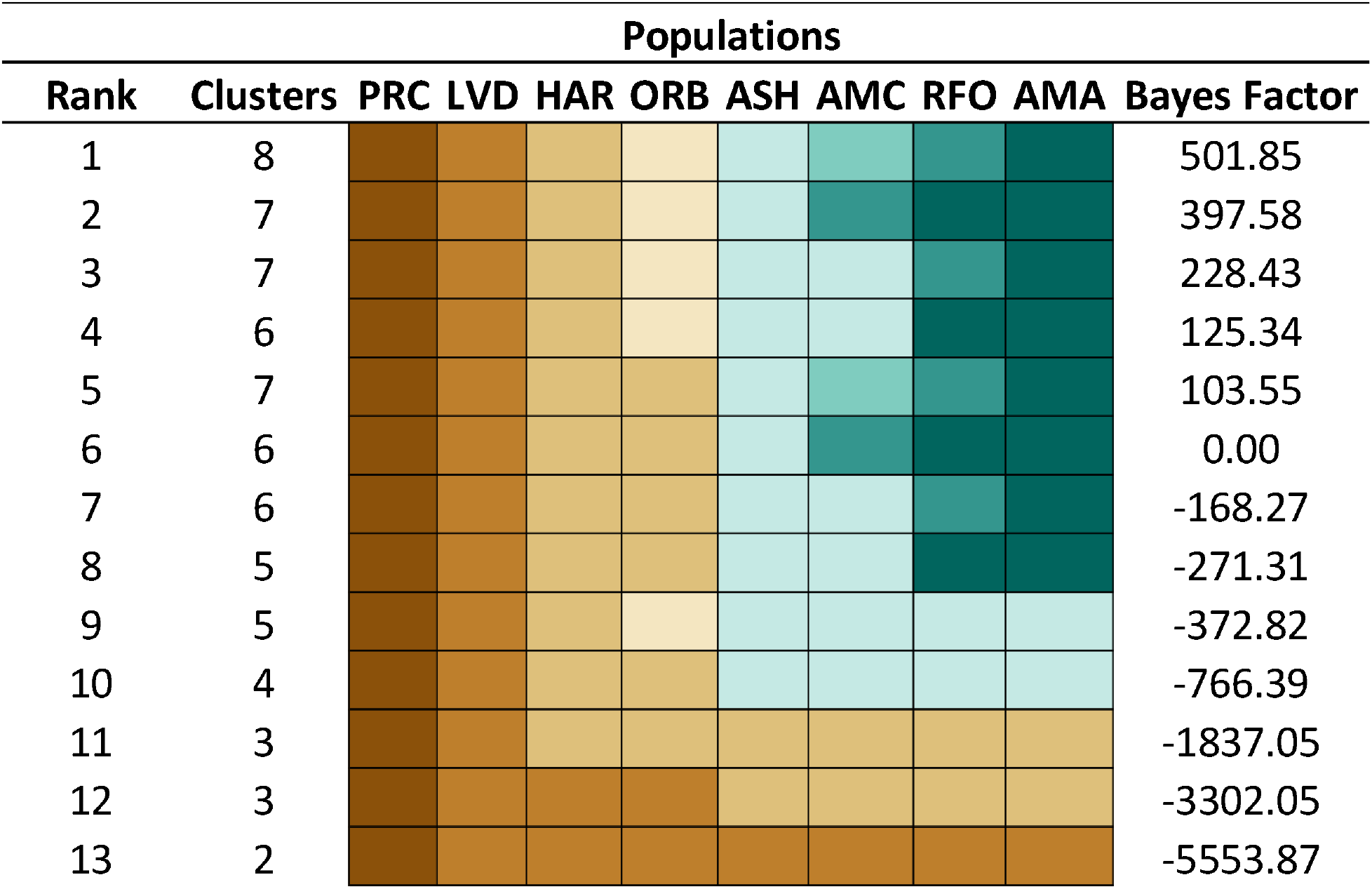
Bayes Factor Delimitation for eight populations of speckled dace (*Rhinichthys osculus*) recovered by Admixture analysis. Five individuals per lineage (N=40) and 200 SNPs were subsampled to satisfy computational restrictions. Thirteen models were tested and ordered by model preference (=Rank) based upon Bayes factors calculated by comparing Marginal Likelihood values for each model computed in SNAPP to the current taxonomy (Bayes Factor=0.00). Models tested a maximum of eight divisions (=Clusters) of populations based upon current distribution and known historical connections among basins. Colors indicate the manner by which locations were grouped for each model (=Rank). SPD lineages are: PRC=Walker Sub-basin (*R. o. robustus*); AMA and RFO=Oasis Valley; AMC=Amargosa Canyon; ASH=Ash Meadows (*R. o. nevadensis*); HAR=Benton Valley; ORB=Owens Valley; LVD=Long Valley.

### 3.6 Unsupervised machine learning

The machine learning methods recovered a minimum of 3 clusters and a maximum of 10 (Figure 7). Random Forest methods, in particular, yielded spurious groupings of individuals that were united by a high proportion of missing data (N=13, 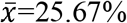, *σ*=17.94%, min=0%, max=53.5%). All other individuals exhibited a much lower proportion of missing data (N=117, 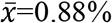, *σ*=2.0%, min=0%, max=16%). Clusters of individuals defined by missing data were typically small in size (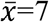 individuals; *σ*=3.39; min=3; max=12).

**Figure 7:**
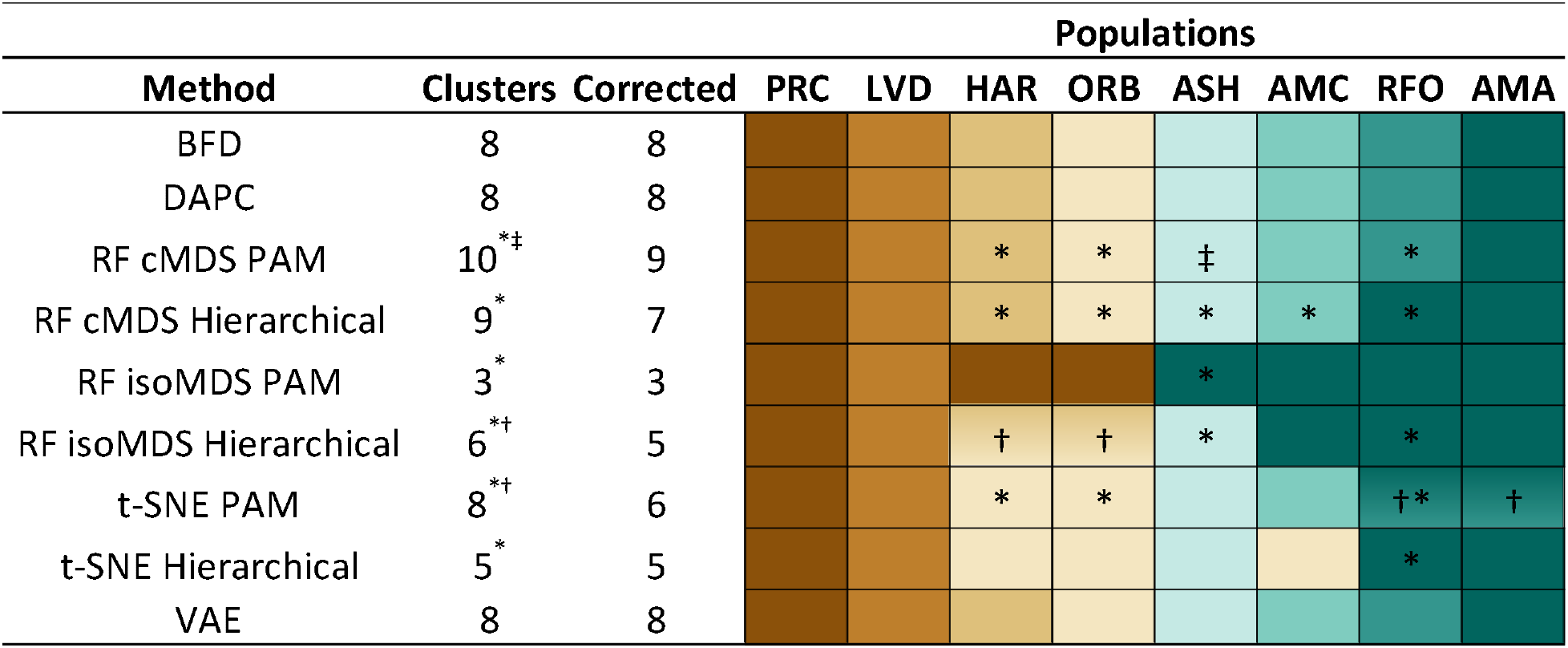
Results of eight unsupervised machine learning (UML) algorithms compared against Bayes Factor Delimitation (BFD*) results. These include four random forest (RF) methods with mixtures of classical (c) and isotonic (iso) multidimensional scaling (MDS) as well as hierarchical and partition around medoids (PAM) clustering. The t-distributed stochastic neighbor embedding (t-SNE) algorithm was employed with two clustering methods. Finally, a discriminant analysis of principal components (DAPC) and variational autoencoder (VAE) were applied. All UML algorithms were applied to 130 individuals genotyped at 200 SNPs. The raw number of divisions (=Clusters) for several methods included groups of individuals that shared a high proportion of missing data (*). Other cases divided individuals from two localities among clusters that did not follow any pattern (†). In one instance, Ash Meadows populations were subdivided according to springs (‡). All three scenarios were interpreted as ‘oversplitting’ and ignored in the final interpretation (=Corrected). SPD lineages are: PRC=Walker Sub-basin (*R. o. robustus*); AMA and RFO=Oasis Valley; AMC=Amargosa Canyon; ASH=Ash Meadows (*R. o. nevadensis*); HAR=Benton Valley; ORB=Owens Valley; LVD=Long Valley.

An unnecessary division of populations occurred in two other instances. The t-SNE PAM algorithm recovered two clusters within Oasis Valley yet neither was clearly defined by a geographic locality (i.e., AMA or RFO). A second instance occurred for Owens Basin localities (HAR and ORB) when evaluated with the RF isoMDS Hierarchical algorithm. Each cluster again contained a mixture of individuals from both localities, so both instances were therefore interpreted as ‘over-splitting.’ Final UML classifications were ‘corrected’ by removing spurious clusters, yielding a minimum of 3 clusters and a maximum of 9.

Results of two algorithms (VAE and DAPC) matched BFD* results. Random Forest and t-SNE methods varied in the number of clusters recovered, with most (5/6; 83.3%) finding fewer clusters than recovered through BFD*. The RF cMDS PAM algorithm was the lone exception. It produced results similar to BFD*, the exception being that nine total clusters were recovered [the ninth a result of Ash Meadows being split according to the two springs sampled in that refuge (ASH and ASR)].

All other RF and t-SNE methods recovered fewer clusters. The RF isoMDS PAM method recovered the fewest (N=3), corresponding to Amargosa Basin (ASH, AMC, RFO and AMA), Long Valley (LVD), and Owens Valley/ Lahontan SPD (HAR, ORB and PRC). The remaining RF and t-SNE methods recovered 5-7 clusters, with Lahontan SPD (PRC), Long Valley (LVD), Ash Meadows (ASH), and Oasis Valley (RFO and AMA) consistently recognized. Discrepancies corresponded to splitting or lumping of Owens Valley populations (HAR, ORB), and the treatment of Amargosa Canyon (AMC) (i.e., unique or lumped with one of the other clusters).

## 4 DISCUSSION

The need for a mechanism to categorize the tips in the tree of life resonates politically, ethically, and biologically (Garnett & Christidis, 2017). Yet, our capacity to do so has languished, not only with regard to the aforementioned species problem, but also with a multiplicity of conservation units at the intraspecific level. The latter represent confusing, and frequently conflicting categorizations that have most often been parsed using phylogeographic analysis (Avise, 2009). However, a more fine-grained phylogenomic perspective is now possible, given the current capacity to generate voluminous data sets encompassing thousands of nuclear loci, and this in turn has subsequently shifted focus and intent (Freudenstein, Broe, Folk, & Sinn, 2017). However, the abilities of researchers to interpret results of phylogenomic studies into discrete conservation units have not kept pace, much to the detriment of adaptive management. A framework is needed to consistently ask if apparent intraspecific boundaries represent factual guides for conservation units, and if so, can they indeed be diagnostically defined (Rannala, 2015)?

One potential mechanism to interpret results of molecular studies that evaluate intraspecific diversity, and to translate them into management solutions would be to place them within a comparative framework with other available data. Here, the difficulty is to (a) know that such data exists, and (b) locate and access it for integrated analyses. In the present situation, this would provide the platform from which results of individual studies could be compiled in a consistent format, compared, and potentially categorized. A recent example is a platform that accommodates a trait-based approach to classification by delineating taxa from which ancestral morphologies and their functions are reconstructed (Gallagher et al., 2020). A similar platform could be built so as to accommodate molecular studies of intraspecific diversity in an attempt to seek consensus amongst results. As a template, it could mirror a recent study (Wieringa et al., 2019) that gathered results of papers published on historical phylogeography in Southeastern North America (N=57), as a mechanism by which new questions relating to intraspecific genetic diversity could be addressed.

### 4.1 The current study

The genetic structure of SPD diagnosed through our analyses reveals their evolutionary history within the DVE. Their distribution is intimately tied to the prehistoric lakes and rivers of the region, with diversifications occurring within modern basins. This pattern clearly reflects a relictual biodiversity with high endemicity, with patterns driven by Pleistocene tectonism and hydrology (i.e., dispersal of SPD from Owens Valley to the Amargosa Basin during fluvial events: Jayko et al., 2008; Knott et al., 2008). However, the current taxonomy is incompatible with these results, and must consequently be adjudicated prior to the delineation of conservation units for management purposes.

In doing so, we first; (A) Associate patterns of population-level diversity with the consensus of species delimitation methods; (B) Address complications resulting from the hybrid status of Amargosa Canyon SPD; (C) Draw comparisons between our results and previous morphological work, as support of intraspecific divisions; and (D) Evaluate genetic variation in the context of previously-recognized subspecific diversity. This allowed us to address the functionality of genomic data as a mechanism for parsing intraspecific biodiversity within both a systematic and conservation context. Finally, we discuss the specific conservation implications regarding the proposed conservation units of SPD in DVE.

### 4.2 Machine learning and subspecific designations

MSC-based species delimitation methods have been successful in delineating taxonomic units within problematic groups (Hedin, 2014; Hedin, Carlson, & Coyle, 2015; Herrera & Shank, 2016). However, caution is a key element in that over-splitting can occur under certain conditions (Sukumaran & Knowles, 2017). Despite these caveats, species delimitation tools remain useful (Leaché et al., 2018) particularly when taxonomic groups are defined according to non-genetic attributes (Barley et al., 2018). In our case, BFD* delimitations correlated with Admixture-defined populations in producing seven DVE lineages.

UML analyses have emerged as an alternative classifier to delineate groups. In comparison to our species delimitation results, UML analyses detected fewer clusters in all instances except two, with a mean of six following adjustments for spurious results due to missing data and over-splitting. The search for a consensus indicated a minimum of four unique DVE conservation units: Long Valley, Owens Valley, Ash Meadows and Oasis Valley (Table 1). We consider Ash Meadows distinct despite it clustering with Oasis Valley in the RF isoMDS PAM results since this algorithm yielded other erroneous groupings that contradict all phylogenetic and population-level analyses (i.e., clustering Owens Valley with *R. o. robustus*).

Our consensus allowed for a relatively robust delimitation of conservation units within the DVE. Despite this, questions still remain with regard to two lineages, Owens and Oasis valleys, in that Admixture indicated differentiated populations within each. Species delimitation methods similarly concurred, with a minority (4/9; 44.4%) splitting Roberts Field (RFO) from the rest of Oasis Valley, whereas a majority (5/9; 55.6%) separated Benton Valley (HAR) from Owens River. The latter is the most intriguing candidate for designation as a separate conservation unit, due largely to its elevated divergence from other local populations, as measured via *F*_ST_. However, a previous phylogenetic reconstruction based on restriction-site mapping of mtDNA failed to distinguish it from Owens River (Oakey et al., 2004), and our study provides moderate support for this same relationship. Unfortunately this population was last sampled in 1989, and may now be extinct as a consequence of flooding that same year (Moyle et al., 2015; Steve Parmenter, California Department of Fish and Wildlife, personal communication).

### 4.3 Hybridization and subspecies

Introgressed populations are another management conundrum (Allendorf, Leary, Spruell, & Wenburg, 2001)(Allendorf et al. 2001) that often have negative consequences for conservation efforts (Rhymer & Simberloff, 1996). However, recent insights from genomic data suggest hybridization is much more common than previously thought (Bangs, Douglas, Brunner, & Douglas, 2020), and in fact reticulation is an important evolutionary process (Chafin, Douglas, Martin, & Douglas, 2019). In our data set, one population, Amargosa Canyon (AMC), appears to be of hybrid origin. Analyses suggest it derived from Admixture between upstream Ash Meadows and Oasis Valley SPD, and is likely a rather recent event (i.e., approximately two generations). This timing estimate may be skewed due to a specific analytical requirement of NewHybrids, where individuals must be classified into pre-defined generational categories. Based upon observed interspecific heterozygosity, our hybrid index contrasting Oasis Valley samples (AMA) with *R. o. nevadensis* (Ash Meadows: ASH) reflects a spread from historical to approximately F2-hybrid status. The HyDe results indicate AMA as being the more probable Oasis Valley parental population than RFO, in that comparisons between the latter and *R. o. nevadensis* were not as well supported. The Amargosa Canyon samples were collected 10-months following a substantial flood that temporarily reconnected this area with upstream populations (i.e., Ash Meadows and Oasis Valley). Interestingly, no pure parentals or F1 offspring were detected, and 10 months is inadequate time for SPD to yield F2 offspring. Thus, we interpret these results as an older hybridization event of indeterminate age.

Given its hybrid origin Amargosa Canyon is our most challenging population to classify. The majority of species delimitation methods (6/9; 66.67%) recognized it as discrete, while those that clustered it with other lineages did so inconsistently. The two RF isoMDS methods clustered Amargosa Canyon with Oasis Valley, while the t-SNE Hierarchical method classified it with Owens River. Neither is consistent with the placement of Amargosa Canyon as sister to *R. o. nevadensis* in the phylogenetic tree. These inconsistencies are likely a reflection of its hybrid ancestry.

### 4.4 Morphological and ecological support for conservation units

Next, we applied a comparative framework to solidify support for our conservation units. While SPD lineages were statistically significant on a genetic basis, they should also be validated morphologically, ecologically, and with additional life history data. This process acts to confirm them as distinct biological entities warranting conservation unit status. A morphological overview of putative subspecies in the DVE focused on broad, regional trends, but it also found “… highly significant differences among all populations for all meristic and mensural characters” (Sada et al., 1995). Ordination also revealed two qualitatively unique body shapes that are typical responses to hydrological conditions (Brinsmead & Fox, 2005; Collin & Fumagalli, 2011). These are: A slender and elongated form typical for flowing streams *versus* shorter and deeper-bodied associated with still water such as lakes or spring. In our study, two populations occur in springs, Long Valley (LVD) and Ash Meadows SPD (ASH/ASR), whereas other study populations are within stream habitats. Benton Valley (HAR) and Owens Valley (ORB) are cold-water streams and irrigation ditches, whereas Oasis Valley (AMA/RFO) and Amargosa Canyon (AMC) are within the Amargosa River. Thus, a contributing factor for morphological and body shape variation is the response by populations to fluvial habitat.

Meristic counts are another type of phenotypic data often applied to diagnose species, but their specificity must be carefully interpreted. Ranges frequently overlap among subspecific divisions, and data for other taxa are either lacking or conflated (Moyle et al., 2015). This is especially true for Oasis Valley SPD, initially lumped with Ash Meadows SPD (Gilbert, 1893; La Rivers, 1962), but with morphological details for subspecific status lacking (Deacon & Williams, 1984; Williams et al., 1982). Likewise, few details are available for Amargosa Canyon SPD (Scoppettone, Hereford, Rissler, Johnson, & Salgado, 2011), other than a series of qualitative descriptors when compared with other SPD subspecies (i.e., smaller head depth, shorter snout-to-nostril length, greater length between anal and caudal fins, greater numbers of pectoral rays, and fewer vertebrae: Moyle et al., 2015). A diagnosis for Ash Meadows SPD is qualitative as well (i.e., incomplete lateral line, relatively large head, small eye, short and deep body, dark stripe along entire length: Gilbert, 1893).

Owens River SPD is also locally variable. Meristic counts (Moyle et al., 2015) are summarized across four populations, to include Benton Valley, thus preventing within-basin comparisons. However, the presence of maxillary barbels distinguishes it from conspecifics in surrounding basins. Furthermore, Benton Valley populations have qualitatively longer pelvic fins, and lower counts for lateral line and pore scales relative to others within-basin (Moyle et al., 2015). In contrast, Long Valley SPD has a higher pectoral and pelvic fin ray count, elevated lateral line scale count, and fewer lateral line pores.

Ecological, life history, and morphological data are thus inconclusive, and fail to delimit conservation units in the DVE. However, this is due to a deficiency of data (rather than homogeneity), and subspecies do seemingly segregate morphologically, albeit without statistical tests as confirmation. In contrast, multiple lines of genetic data provide clear signals of distinct entities, and inferences from modern genomic techniques also reinforce observed gaps in ecological data (Crandall, Bininda-Emonds, Mace, & Wayne, 2000; Funk, McKay, Hohenlohe, & Allendorf, 2012). *F*_ST_ outlier loci, for example, diagnosed potential ecological adaptation among SPD lineages, in that loci under selection have the potential to reveal cryptic signals of adaptive divergence (Tigano, Shultz, Edwards, Robertson, & Friesen, 2017). While these loci do not replace traditional field observations (Funk et al., 2012), they do underscore in our situation the juxtaposition of neutral variation and adaptive variation among DVE units. Isolation in different habitat types (i.e., springs *versus* rivers) therefore highlights the need for conservation strategies that differ yet are linked via an ecosystem-oriented focus.

### 4.5 Conservation implications with regard to subspecies

Modern genomic tools offer an in-depth view of population histories and allow unique genetic lineages to be discriminated. In our study, they phylogenetically validate a rare situation where the majority (80%) of anecdotal subspecific designations (our OTUs) are recognized as genetically discrete units. We deem this as a response to the extreme geographic isolation imposed upon each DVE lineage. Until formal descriptions can occur (i.e., morphological analyses so as to formally describe each lineage), we thus recognize Long Valley SPD, Owens Valley SPD, and Oasis Valley SPD as ESUs (Moritz, 1994; Ryder, 1986; Waples, 1991), and substantiate Ash Meadows SPD (*R. o. nevadensis*) as a distinct taxon and conservation unit.

Endemicity resulting from habitat isolation provides a unique challenge for conservation. While isolated habitats are often prone to human disturbance, they are readily identified and monitored as well (Arthington, Dulvy, Gladstone, & Winfield, 2016). Springs, for example, exhibit high levels of genetic structure across small geographic areas (Echelle et al., 2015). They frequently reflect low within-system species richness (α-diversity) but high diversity when compared to other systems (β-diversity) (Gibert et al., 2009). In other words, greater diversity exists between groundwater-dependent systems rather than within (Gibert & Deharveng, 2002). Thus, groundwater depletion invokes dire consequences for aquatic fauna, ranging from the depletion of faunal assemblages (Perkin et al., 2017) to complete extinction (Miller, Williams, & Williams, 1989). The latter scenario provides an elevated risk for these populations since they lack redundancy to protect from catastrophic loss.

Our validation of anecdotal subspecies has a conservation imperative, in that it underscores the necessity of a management strategy that sustains and protects these unique entities. Practical considerations require an adherence to current conservation policy, and this remains species-centric (Smith et al., 2018). While it has benefits (Runge et al., 2019), ecosystem-level factors are also necessary conservation aspects (Franklin, 1993). They are particularly important in regions containing multiple narrowly endemic species, as those rare and sympatric can suffer unintentional consequences through a strict species-centric approach (Casazza et al., 2016). Importantly, climate change impacts are also relevant for ecosystem conservation (Prober, Doerr, Broadhurst, Williams, & Dickson, 2019; Wilkening, Pearson-Prestera, Mungi, & Bhattacharyya, 2019).

In addition, the evolutionary history of these lineages adds complications. In the DVE, each lineage of SPD is narrowly endemic and scant evidence of contact zones, with the exception of the hybrid population in Amargosa Canyon (AMC). Hydrological changes underlying these patterns occurred on considerably different timescales in the two basins. Loss of surface water in the Owens Basin is directly tied to anthropogenic activities during the past century, with available habitat being severely restricted (Buckmaster & Parmenter, 2019). The full extent of how much this depleted genetic diversity is unknown, but our data document the probable contemporary loss of (at least) one genetically distinct lineage from this system (i.e., RFO in the Oasis Valley).

Finally, genomic tools have demonstrated the previously unknown hybrid status of Amargosa Canyon SPD. Hybridization has been commonplace among desert fishes (Bangs, Douglas, Mussmann, & Douglas, 2018) and has served as a mechanism of speciation (Gerber, Tibbets, & Dowling, 2001), but can also erode species boundaries (Chafin, Douglas, Martin, & Douglas, 2019). Additionally, anthropogenic climate change has induced hybridization among divergent species (Canestrelli et al., 2017; Muhlfeld et al., 2014), and thus represents a post-Pleistocene evolutionary mechanism inherent to western North America (Woodhouse, Meko, MacDonald, Stahle, & Cook, 2010).

Hybridization is a contentious conservation topic (Fitzpatrick, Ryan, Johnson, Corush, & Carter, 2015), particularly when one parental species is afforded protection under the US Endangered Species Act (ESA), as is the case with Ash Meadows SPD (*R. o. nevadensis*). The issue is further compounded by a lack of policy explicitly addressing hybrids (vonHoldt, Brzeski, Wilcove, & Rutledge, 2017). The historic and ongoing lineage mixing that yielded Amargosa Canyon SPD (AMC) appears natural and thus should not preclude protection (Allendorf et al., 2001). The situation parallels that of the red wolf (*Canis rufus*), deemed a hybrid between endangered grey wolf (*Canis lupus*) and coyote (*Canis latrans*) (Hohenlohe et al., 2017; vonHoldt et al., 2016), although recent genomic analyses revealed it to be a distinct species of its own with a convoluted history of introgression (Chafin, Douglas, & Douglas, 2020). Given this precedence, Amargosa Canyon SPD could therefore be listed as a DPS under the ESA (Waples, Kays, Fredrickson, Pacifici, & Mills, 2018).

## 5 CONCLUSIONS

Distinct SPD lineages within the DVE represent Pleistocene hydrological connections among basins. They are narrowly endemic and relictual components of a more pluvial Pleistocene ecosystem that now persists as small pockets within desert oases. The majority represent entities previously identified anecdotally, yet without formal description. Despite their academic recognition, legal protection is absent and their existence remains manifestly precarious, save for one described entity (*R. o. nevadensis*). Our results demonstrate the necessity of using multiple approaches in a comparative framework to diagnose conservation units (Figure 1). They also add to the growing body of literature that indicates MSC-based species delimitation methods serve to demarcate populations rather than species. Our results sustain one subspecies (*R. o. nevadensis*), validate three lineages (Oasis Valley, Owens River, and Long Valley) as distinct ESUs, and argue that Amargosa Canyon SPD is eligible for protection as a DPS under existing environmental laws.

## ACKNOWLEDGMENTS

This research represents partial fulfillment of the Ph.D. degree (SMM) in Biological Sciences at the University of Arkansas. It was supported by generous University endowments: The Bruker Professorship in Life Sciences (MRD), the 21^st^ Century Chair in Global Change Biology (MED), and a Doctoral Academy Fellowship (SMM). Computational resources were provided by the Arkansas High Performance Computing Center (AHPCC) and the NSF Jetstream XSEDE Resource (XSEDE Allocation: TG-BIO160065). Sampling procedures were approved under Arizona State University Animal Care and Use Committee (ASU IACUC) permit 92-0242R. Several state and federal agencies contributed permit approvals, field expertise, samples, and valuable comments: California Department of Fish and Wildlife, Nevada Department of Wildlife, and the United States Fish & Wildlife Service (permits PRT-676811, PRT-797129, and TE-702631). In particular, we thank Matt Andersen, Alejandra Cortes, Shawn Goodchild, Kevin Guadalupe, James Harter, Greg Munson, Steve Parmenter, and Donald Sada. Several members of the Douglas Lab contributed time to DNA work over the years, including Max Bangs, Tyler Chafin, Jennifer Cotter, and Alex Klug. The use of trade, product, industry or firm names is for informative purposes only and does not constitute an endorsement by the U.S. Government or the U.S. Fish and Wildlife Service. Links to non-Service Web sites do not imply any official U.S. Fish and Wildlife Service endorsement of the opinions or ideas expressed therein or guarantee the validity of the information provided. The findings, conclusions, and opinions expressed in this article represent those of the authors, and do not necessarily represent the views of the U.S. Fish & Wildlife Service.

## CONFLICT OF INTEREST

None Declared.

## AUTHOR CONTRIBUTIONS

All authors participated in study design; SMM prepared DNA, generated ddRAD libraries, and completed data analysis; all authors contributed in drafting the manuscript and all approved its final version.

## DATA ACCESSIBILITY

Raw fastq files for each individual as well as all alignments used in this study are available on the Dryad Digital Repository (address added after acceptance of article).

## Appendix 1 Death Valley Subspecies

Speckled dace within the Death Valley region exhibit high levels of diversity among the many isolated sites they inhabit. Population isolation has led to the proposal of five SPD subspecies within the Owens (N=2) and Amargosa (N=3) watersheds. All five subspecies are recognized by Nevada and California as being species of special concern (Moyle *et al.* 2015).

Two subspecies are found in the Owens River Valley: Long Valley SPD (*R. o.* ssp 12), and Owens River SPD (*R. o.* ssp 2). Long Valley subspecies is the most imperiled freshwater fish in California (Moyle *et al.* 2011). It is feared extirpated from the wild as of 2019. Its sole habitat is restricted to Whitmore Hot Springs - a thermal-spring complex that receives the partially chlorinated outflow of a public swimming pool (Moyle *et al.* 2015). Both morphological and genetic studies have confirmed its status as a distinct taxon, but to date it has not been formally described (Sada *et al.* 1995; Oakey *et al.* 2004; Furiness 2012).

The Owens River SPD also has a tenuous existence. The subspecies has been extirpated from eight of 17 historically known sites, and existing populations are only left in Fish Slough, Round Valley, and areas around Bishop, CA (Moyle *et al.* 2015). Habitat fragmentation has occurred due to export of water to Los Angeles following construction of aqueducts in 1913 and 1970 (Hollett *et al.* 1991). This subspecies was once grouped with Ash Meadows SPD (Gilbert 1893) until being recognized as a distinct population (Williams *et al.* 1982; Deacon & Williams 1984). Sada et al. (1995) also recognized morphological and genetic variation among Owens River populations, but did not determine additional splitting of the subspecies was warranted.

Three additional subspecies are found in the Amargosa Drainage: Ash Meadows SPD (*R. o. nevadensis*), Amargosa Canyon SPD (*R. o.* ssp 1), and Oasis Valley SPD (*R. o.* ssp 6). Ash Meadows SPD is the only species in the region afforded federal protection under the U.S. Endangered Species Act (ESA: Federal Register 1983). This subspecies is found within Ash Meadows National Wildlife Refuge, which was founded in 1984 to provide protection for all of the unique fauna endemic to Ash Meadows (Deacon & Williams 1991). This taxon was initially described as a different species (*R. nevadensis*: Gilbert 1893) before later being designated a subspecies of *R. osculus* (Hubbs *et al.* 1974).

The upper Amargosa River is home to Oasis Valley SPD. It is currently restricted to Fleur de Lis Springs and a short portion of the upper Amargosa River near Beatty Nevada (Sada *et al.* 1995). Amargosa Canyon SPD is found in the lower Amargosa River, and has been evaluated to be in danger of extinction within the next 50 years (Moyle *et al.* 2011). Today it is mostly confined to Willow Creek and Willow Creek Reservoir (Williams *et al.* 1982; Scoppettone *et al.* 2011). It has been extirpated from a spring north of Tecopa (Miller 1938; Moyle *et al.* 2015). This subspecies was once grouped with *R. o. nevadensis* (Gilbert 1893; La Rivers 1962) and later recognized as distinct from the Ash Meadows subspecies (Williams *et al.* 1982; Deacon & Williams 1984).

## Notes

### Competing Interest Statement

The authors have declared no competing interest.

